# A cell-free system for functional studies of small membrane proteins

**DOI:** 10.1101/2023.12.22.573026

**Authors:** Shan Jiang, Gülce Çelen, Timo Glatter, Henrike Niederholtmeyer, Jing Yuan

**Affiliations:** Max Planck Institute for Terrestrial Microbiology and Center for Synthetic Microbiology, 35043 Marburg, Germany; Technical University of Munich, Campus Straubing for Biotechnology and Sustainability, 94315 Straubing, Germany

**Keywords:** Small membrane proteins, cell-free synthesis, lipid sponge droplets

## Abstract

Numerous small proteins have been discovered across all domains of life, among which many are hydrophobic and predicted to localize to the cell membrane. Based on a few that are well-studied, small membrane proteins are regulators involved in various biological processes, such as cell signaling, nutrient transport, drug resistance, and stress response. However, the function of most identified small membrane proteins remains elusive. Their small size and hydrophobicity make protein production challenging, hindering function discovery. Here, we combined a cell-free system with lipid sponge droplets and synthesized small membrane proteins *in vitro*. Lipid sponge droplets contain a dense network of lipid bilayers, which accommodates and extracts newly synthesized small membrane proteins from the aqueous surroundings. Using small bacterial membrane proteins MgrB, SafA, and AcrZ as proof of principle, we showed that the *in vitro* produced membrane proteins were functionally active, for example, modulating the activity of their target kinase as expected. The cell-free system produced small membrane proteins, including one from human, up to micromolar concentrations, indicating its high level of versatility and productivity. Furthermore, AcrZ produced in this system was used successfully for *in vitro* co-immunoprecipitations to identify interaction partners. This work presents a robust alternative approach for producing small membrane proteins, which opens a door to their function discovery in different domains of life.

**Importance:** Small membrane proteins are shown to be involved in various biological processes in all domains of life and “can no longer be ignored”. Due to their small size and hydrophobicity, functional investigation of small membrane proteins is challenging. In this work, we present a simple, versatile, cell-free approach for synthesizing small membrane proteins *in vitro*. We show that the small membrane proteins produced with our system are functional and in sufficient amounts for downstream target discoveries. Furthermore, our approach may uncover additional regulatory functions of small membrane proteins studied with conventional methods. Our work provides a robust alternative workflow for functional studies, which opens up new possibilities to advance our understanding of small membrane protein biology.

## Introduction

Small proteins also named mini-proteins [1], micro-proteins [2], or small peptides [3], were overlooked in the past due to their small size (less than 50 amino acids in prokaryotes and less than 100 amino acids in eukaryotes) [3-5]. Small proteins are directly translated from small open reading frames (sORFs) that often start with noncanonical start codons, which increases the difficulty of differentiating valid sORFs from noncoding RNAs [3, 4]. With the advances in bioinformatics and application of ribosomal profiling, hundreds to thousands of sORFs have been identified in recent years [6-8]. While some have low stability and have no apparent contribution to cell fitness in the conditions tested [9], many are conserved, predicted to have a putative transmembrane helix, and may be involved in cell signaling and communication [6, 10].

Small membrane proteins constitute a significant portion of newly discovered small proteins. They are found to be involved in diverse biological processes across domains of life, ranging from bacterial virulence regulation to immunosurveillance in mammalian cells (reviewed in [11, 12]). Well-characterized bacterial small membrane proteins primarily act as regulators by integrating into large protein complexes, such as KdpF facilitating potassium transport complex formation [13], AcrZ specifying substrates for the AcrBA-TolC drug efflux pump [14], and Prli42 anchoring the stressosome to the bacterial membrane [15]. Small membrane proteins also target individual larger membrane proteins, regulating their stability and function. For example, MgrB and SafA participate in cell signaling by regulating the activity of the sensor kinase PhoQ [16-19]. MgtS and MntS regulate magnesium and manganese transport by targeting the respective transporter proteins [20, 21]. Lastly, some small membrane proteins directly target the cell membrane, acting as a detector for membrane curvature [22] or as type I toxins regulating membrane permeability [23].

Despite the knowledge and experience gained from the well-characterized small proteins, the function of most newly discovered small membrane proteins awaits further investigation. In contrast to soluble proteins, small membrane proteins are challenging to produce via chemical synthesis due to their hydrophobicity. Overexpression of recombinant small membrane proteins *in vivo* may induce general stress or even be toxic to cells. These difficulties become major roadblocks for functional studies.

Cell-free protein synthesis (CFPS) has been used for membrane protein production *in vitro* to avoid potential cytotoxicity and protein aggregation, to improve yield, and to characterize membrane protein functions [24-27]. Commonly used hydrophobic supplements in a cell-free system for membrane protein production include detergents, nanodiscs, and liposomes, which prevent precipitation and support the synthesis and folding of membrane proteins during synthesis. Liposomes closely mimic cellular membranes. Their membrane surface area correlates with the yield of folded and active membrane proteins [28, 29], underscoring the importance of a suitable membrane environment for functional membrane protein production in CFPS. Liposomes also generate separate compartments, which are required for functional assessment of channel and transporter proteins. However, the main drawback of liposomes is their instability in time and response to osmolarity changes. Nanodiscs [30] are a more stable alternative to liposomes, but both require specialized equipment and multiple steps for their preparation. In addition, both systems are limited in the membrane surface area they can provide.

Unlike liposomes and nanodiscs, lipid sponge droplets contain dense non-lamellar lipid bilayer networks, providing an extensive membrane surface for membrane protein insertion. They have a high water content, harbor nanometric aqueous channels, and thus simultaneously accommodate hydrophilic components of a cell-free system. Lipid sponge droplets are simple to prepare and generated via spontaneous assembly of the single-chain galactolipid N-oleoyl β-D-galactopyranosylamine (GOA) and non-ionic detergents such as octylphenoxypolyethoxyethanol (IGEPAL). In aqueous solutions, the two amphiphiles form micrometer-sized coacervate-like droplets that are stable in a wide range of conditions [31, 32]. Despite being a nonbiological lipid, the thickness of GOA membranes (35.25 Å)[33] is comparable to that of a biological membrane (37-40 Å) [34], which may explain why membrane proteins such as diacylglycerol kinase and cytochrome c oxidase were functional when reconstituted in lipid sponge droplets [31].

In this study, we combined lipid sponge droplets with a minimal, defined cell-free system (PURE system) [35] and produced small membrane proteins *in vitro*. We chose the PURE system because of its compatibility with the droplets and to avoid potential interference with downstream functional studies of the synthesized small membrane proteins. Using *E. coli* proteins MgrB and SafA as proof of principle, we showed that the small proteins were successfully synthesized and localized in the lipid sponge droplets. The yield was sufficient for downstream functional assays, and the lipid sponge was required to reach this level of protein production. The synthesized MgrB and SafA were functionally active, modifying the kinase activity of their target protein PhoQ *in vitro*. Successful synthesis of two other small membrane proteins AcrZ from *E. coli* and sarcolipin from human, demonstrates the versatility of this cell-free system, and we present strategies to increase the yield of the synthesized small membrane proteins to micromolar concentrations. Lastly, we successfully co-immunoprecipitated the interacting proteins of AcrZ from solubilized cell membranes, suggesting that this cell-free system is suitable for target discovery of small membrane proteins forming stable complexes with their targets.

## Results

### Cell-free biosynthesis of small membrane proteins MgrB and SafA

We started with two well-studied *E. coli* proteins MgrB (47aa) and SafA (65aa) to test the cell-free system for small membrane protein production. DNA templates containing T7 promoter and the ORFs of MgrB and SafA were constructed (**Fig. 1**) and used to initiate protein synthesis in the cell-free system with lipid sponge droplets. To visualize CFPS, we added an N-terminal mNeonGreen tag to the small membrane proteins. Additionally, a DNA template for synthesizing only mNeonGreen was used as a control. We observed a gradual increase in green fluorescence during incubation, which reached a plateau after around three hours (**Fig. 1, Movie S1**). For the synthesis of mNeongreen-MgrB and mNeongreen-SafA, the fluorescence primarily localized to lipid sponge droplets (**Fig. 1, Movie S1**). Conversely, the mNeongreen control sample showed diffuse fluorescence in the field of view with lipid sponge droplets having a slightly lower fluorescent intensity (**Fig. 1C**). These results suggested that the mNeongreen-tagged small membrane proteins were produced in the cell-free system and that their hydrophobicity led to sequestration of the fusion proteins into the membrane-rich droplet phase. To quantify MgrB and SafA synthesized in the cell-free system, we separately introduced a FLAG epitope tag to the N-terminus of the proteins (**Fig. 2A**). Using a commercially available FLAG-tagged protein as a reference in Western blot analysis, we quantified that approximately 600 nM FLAG-SafA was produced in the cell-free system after three hours and about 300 nM for FLAG-MgrB (**Fig. 2B, Fig. S1**). This concentration is comparable (in terms of mg/ml) to the reported concentrations reached for larger membrane proteins in PURE reactions supplemented with liposomes [36].

**Fig. 1.**
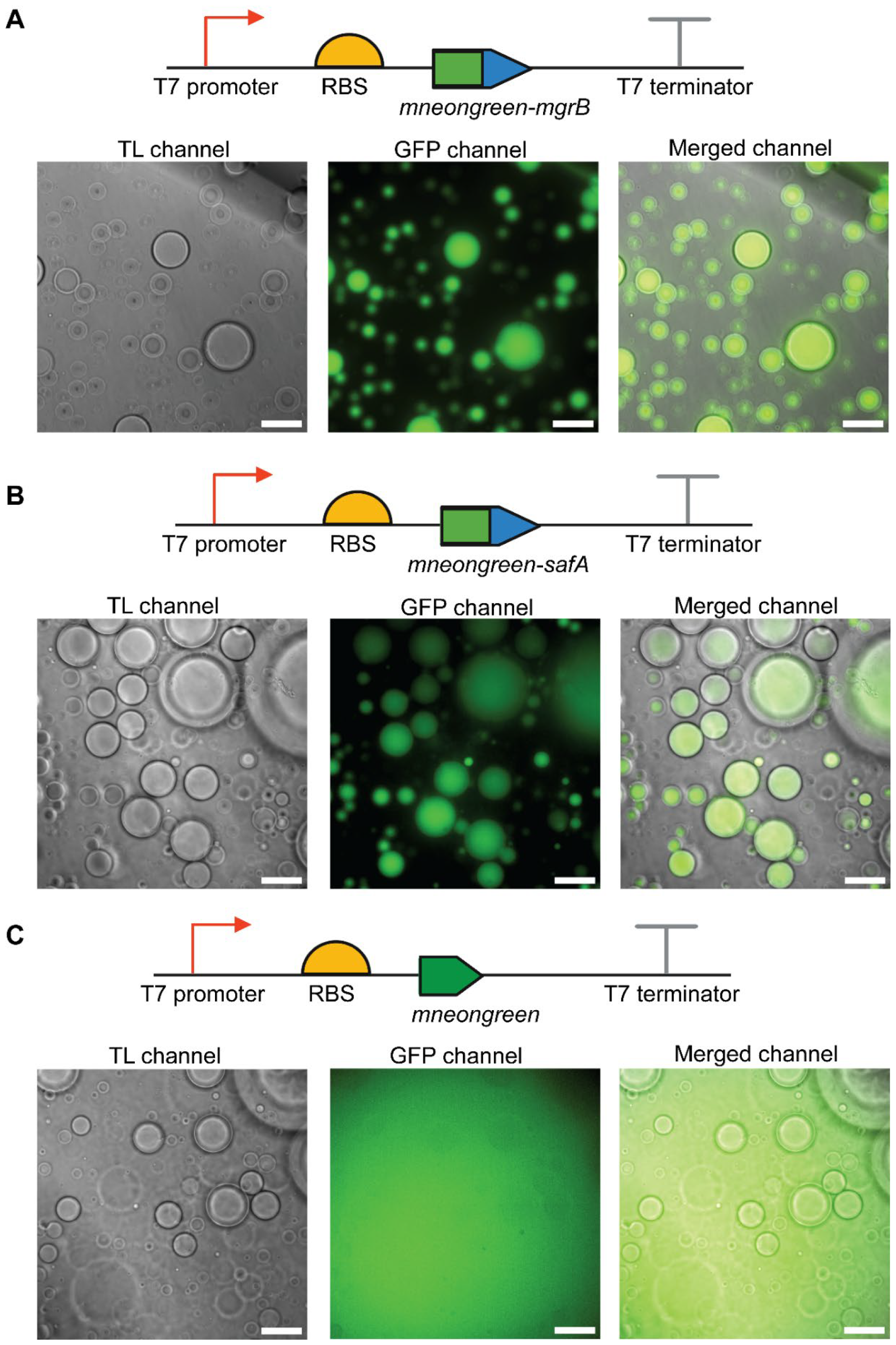
Cell-free synthesis of small membrane proteins in the presence of lipid sponge droplet. The mNeonGreen labeled MgrB (A), SafA (B), and mNeonGreen control (C) were synthesized in the cell-free system in the presence of lipid droplets. Representative images from the transmitted light (TL), GFP fluorescence and merged channels are shown with the corresponding DNA template scheme above (Scale bar, 50 μm). Data are representative of at least three independent experiments.

**Fig. 2.**
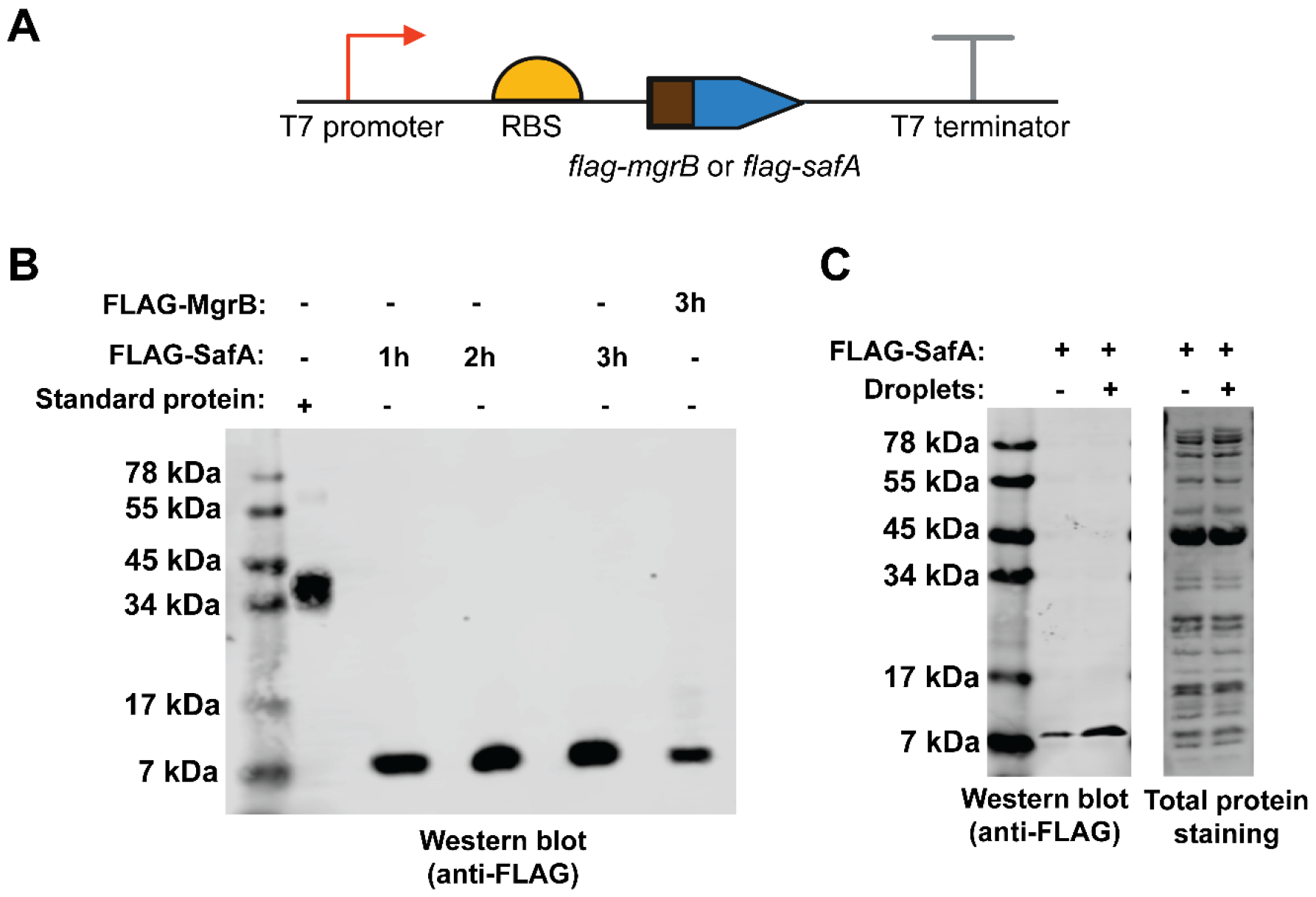
Quantification of synthesized small membrane proteins in the cell-free system. **(A)** Design of the linear DNA templates. **(B)** Western blot analysis of the synthesized MgrB and SafA in a 5 μL reaction. A FLAG-tagged standard protein (0.1 μg) was used as a positive control. **(C)** Cell-free synthesis of SafA with or without lipid sponge droplets. Data are representative of three independent experiments.

Small membrane proteins, such as MgrB and SafA, typically contain a single transmembrane helix (**Table S1**). To determine whether lipid sponge droplets were required for such bitopic integral membrane protein synthesis *in vitro*, we produced and compared FLAG-SafA with the PURE system in the presence and absence of lipid sponge droplets. The Western blot results showed that a higher amount (2.73 times) of SafA was produced in the presence of lipid sponge droplets (**Fig. 2C, Fig. S2**), indicating that the hydrophobic environment is beneficial despite the seemingly simple tertiary structure of SafA. The lipid sponge droplets facilitated the reconstitution of the synthesized protein, likely prevented protein aggregation, and thus increased the yield of SafA during CFPS. Along similar lines, previous work on CFPS of membrane channel proteins showed that the production of folded and active protein correlated with the amount of supplemented liposomes [28, 29]. While the membrane surface area liposomes can offer is limited, lipid sponge droplets are filled with a dense membrane environment and can be added to CFPS reactions at high concentrations [31].

### Functional analysis of the synthesized MgrB and SafA

The function of MgrB and SafA can be tested by following the activity of their target protein PhoQ. The sensor kinase PhoQ catalyzes an autophosphorylation reaction, where it utilizes ATP to phosphorylate a conserved histidine residue. MgrB and SafA directly interact with PhoQ. MgrB inhibits the autophosphorylation of PhoQ, while SafA is an activator. To test the regulatory function of the synthesized small proteins, we performed PhoQ autophosphorylation assays with mNeonGreen as a negative control (the experimental scheme shown in **Fig. 3A**). To introduce the synthesized FLAG-SafA and FLAG-MgrB to the reaction, we spun down the droplets after CFPS and replaced the supernatant with purified recombinant PhoQ protein in phosphorylation buffer. Notably, residual aqueous components of the cell-free system were carried over to the autophosphorylation reaction due to the unique three-dimensional structure of lipid sponge droplets. The reaction mix was divided into two aliquots. One proceeded to autophosphorylation assays by adding radioactively labeled [1-^32^P]ATP. The other was used for SDS-PAGE/Western blot analysis to verify the production of the small proteins and the amount of PhoQ. We observed an increase in radioactively labeled PhoQ in the presence of the synthesized FLAG-SafA and a decrease in autophosphorylation with the synthesized FLAG-MgrB compared to the negative control (**Fig. 3B, 3C**), indicating that the synthesized small membrane proteins are functionally active.

**Fig. 3.**
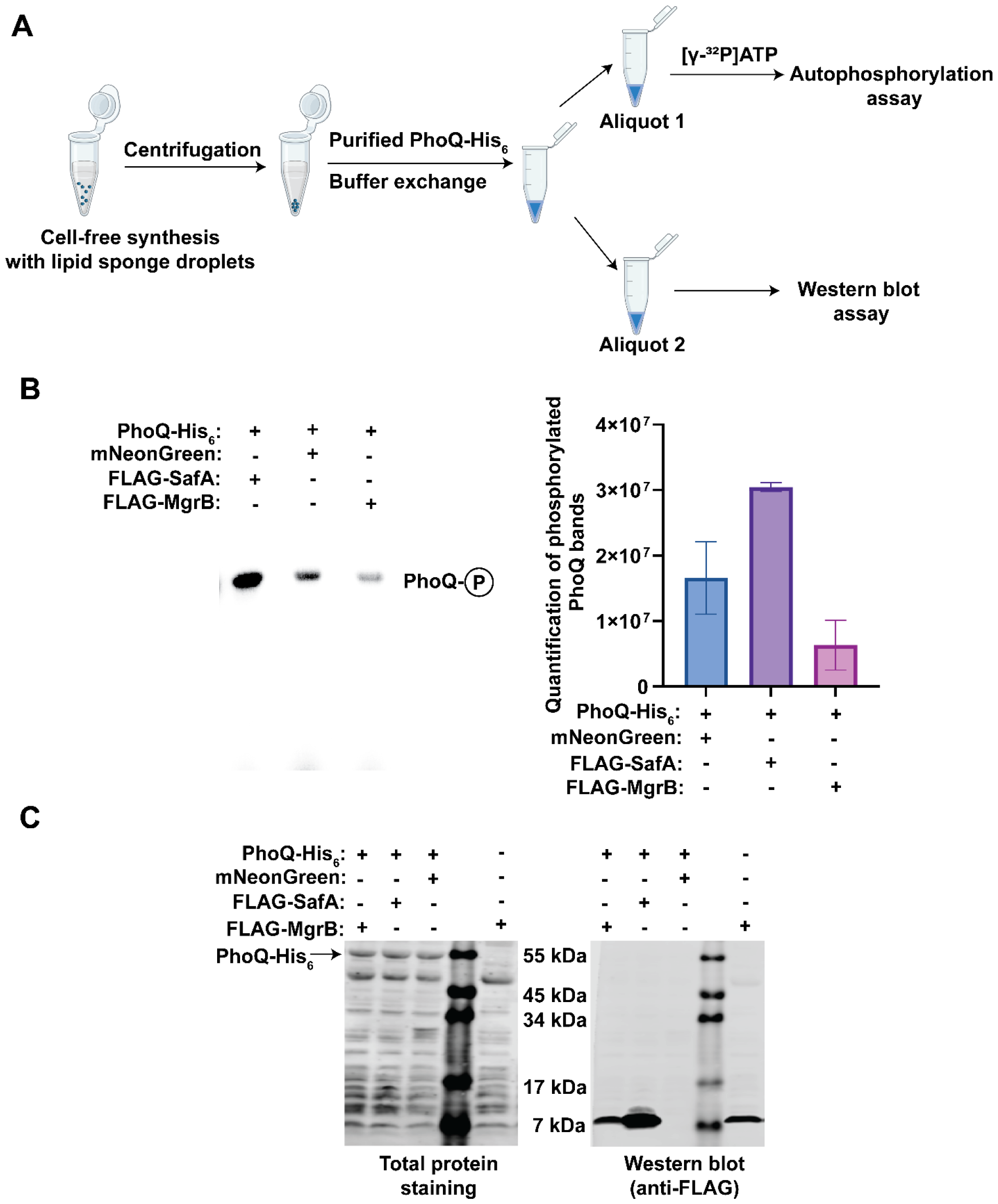
Functional assay of synthesized small membrane proteins MgrB and SafA. **(A)** Scheme of the functional assay. **(B)** Autophosphorylation of PhoQ in the presence of synthesized proteins. The reactions were carried out at room temperature for 30 minutes. The phosphorylated PhoQ was separated from free [γ-^32^P]ATPs by SDS-PAGE and detected with a phosphorimager (left). The bands were quantified using ImageJ software (right). Error bars represent the standard deviations from three independent experiments. **(C)** The total protein and Western blot analysis of the reaction mix in the functional assay. Data are representative of at least three independent experiments.

With the same setup, we synthesized and tested the function of MgrB and SafA without epitope tags, and observed similar results with autophosphorylation assays (**Fig. S3**). Altogether, the results suggested that our cell-free system can produce functional small membrane proteins in sufficient amounts for downstream analysis *in vitro*. The ability to synthesize native functional small membrane proteins is especially advantageous for studying the ones sensitive to epitope or affinity tags.

### Cell-free synthesis of other small membrane proteins

To test the versatility of the cell-free system, we applied it to synthesize two other small membrane proteins, AcrZ from *E.coli* and sarcolipin from human. AcrZ is an integral membrane protein. It interacts with the drug efflux protein AcrB and forms a 3:3 hexameric membrane protein complex, which can bind to AcrA and TolC, forming a complete drug efflux pump [14, 37]. Functionally, AcrZ regulates the substrate specificity of the drug efflux transporter [14, 37]. Sarcolipin is an essential regulator of the sarco-/endoplasmic reticulum Ca^2+^-ATPase in human myocytes [38]. It prevents the pumping of calcium cations out of the cytosol without inhibiting the ATPase activity, thus converting the chemical energy stored in ATP to heat [38]. After codon optimization, we constructed the DNA templates and performed CFPS of AcrZ and sarcolipin in the presence of lipid sponge droplets. A FLAG-tag was introduced for quantification via Western blot analysis. With SafA as a positive control, we observed that AcrZ and sarcolipin were produced using the cell-free system. The yield of AcrZ was about 2.4 times more than SafA, reaching the micromolar range (about 1.4 µM) (**Fig. 4A, Fig. S4**). Human sarcolipin was synthesized less (about 90 nM). One reason for the lower yield could be the polar residue (Asn11) in its transmembrane helix (**Table S1, Fig. 4B, Fig. S5**) hindering protein partitioning to lipid sponge droplets. Indeed, an N to L substitution increased the yield (approximately 2.4 times) of sarcolipin (**Fig. S6**).

**Fig. 4.**
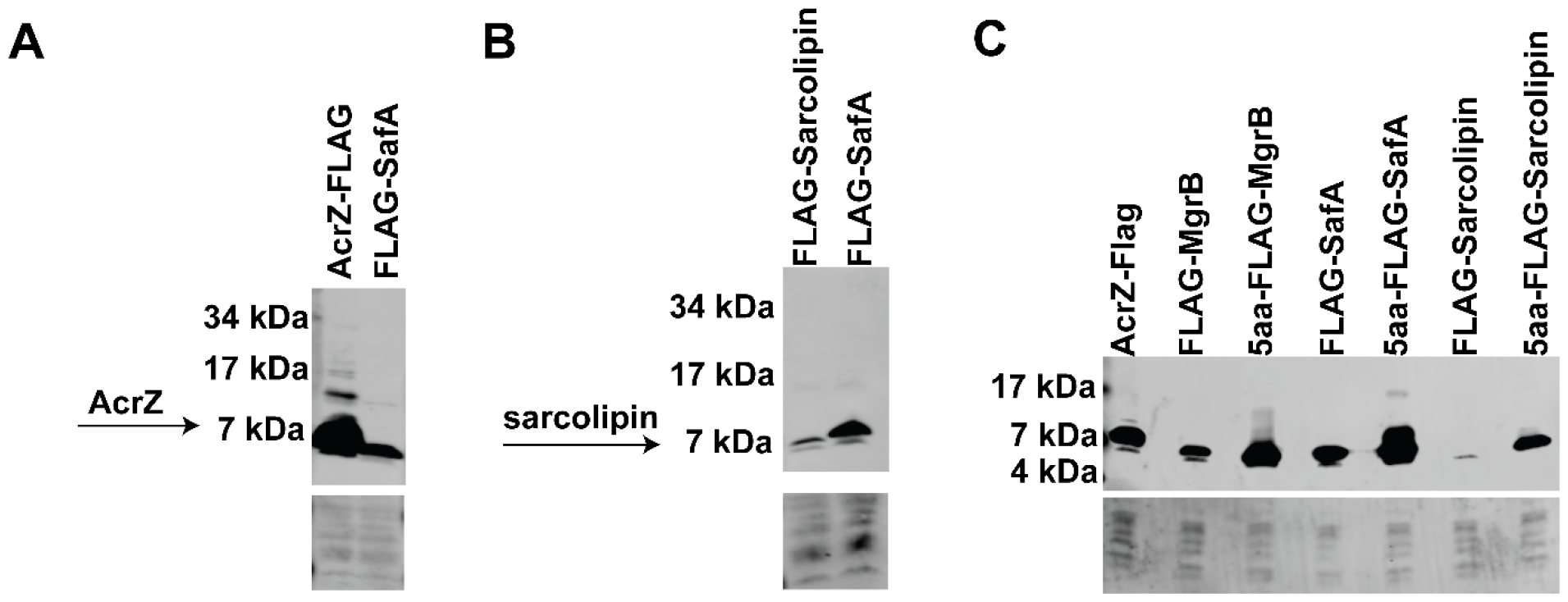
Cell-free synthesis of AcrZ, sarcolipin, and yield optimization. Western blot analysis of the synthesized FLAG-tagged AcrZ from *E. coli* **(A)** and sarcolipin from human **(B)** are shown with the synthesis of SafA as a positive control. **(C)** Western blot analysis of synthesized small membrane proteins with or without N-terminal 5-amino acid (5aa) insertions. The corresponding total protein stain of the PVDF membranes serves as loading control. Data are representative of three independent experiments.

A previous study showed that the mRNA sequence immediately downstream of the start codon affects translation efficiency [39]. To increase the yield of MgrB, SafA, and sarcolipin to a level comparable to that of AcrZ, we then copied the DNA sequence coding for the first five amino acids of AcrZ and inserted it between the start codon and the N-terminal FLAG tag in the corresponding ORFs. When using these modified DNA templates for cell-free synthesis, we observed a drastic increase in protein yield (**Fig. 4C**). All three small membrane proteins were produced near or at micromolar concentrations (**Fig. S7**), significantly more than the previously reported concentrations for membrane protein synthesis in PURE reactions with liposomes [36]. Taken together, our results demonstrate the versatility and productivity of the cell-free system in generating small membrane proteins.

### Target discovery using the synthesized small membrane proteins

To determine whether the *in vitro* synthesized small membrane proteins were suitable for target discovery, we used MgrB, SafA, and AcrZ fused to a FLAG tag as bait to identify their interaction partners via co-immunoprecipitation (co-IP) (**Fig. 5A**). The tagged small membrane proteins were synthesized using the cell-free system as described above. Meanwhile, we isolated the total membrane fraction from the corresponding *E. coli* knockout strains (*ΔmgrB, ΔsafA*, and *ΔacrZ*). The membrane was solubilized with a detergent-containing buffer and incubated with the synthesized FLAG-tagged small membrane proteins. In this step, the detergent dissolved most lipid sponge droplets, allowing interactions between the synthesized small protein and components in the cell membrane. With anti-FLAG magnetic beads, proteins bound to bait were analyzed with mass spectrometry (MS), and the results showed that seven proteins were reproducibly enriched in the AcrZ co-IP samples with Z-scores above 2 (**Fig. 5B, File S1 and S2**). AcrB, the known interactor of AcrZ, was identified as the top hit with the highest spectrum counts. AcrA, a periplasmic protein that can bind to AcrB and TolC, forming a tripartite multidrug efflux pump, was also enriched with fewer spectrum counts. The other five enriched proteins (MdtF, ArcB, Trg, MrcA, and LptD) are all membrane proteins.

**Fig. 5.**
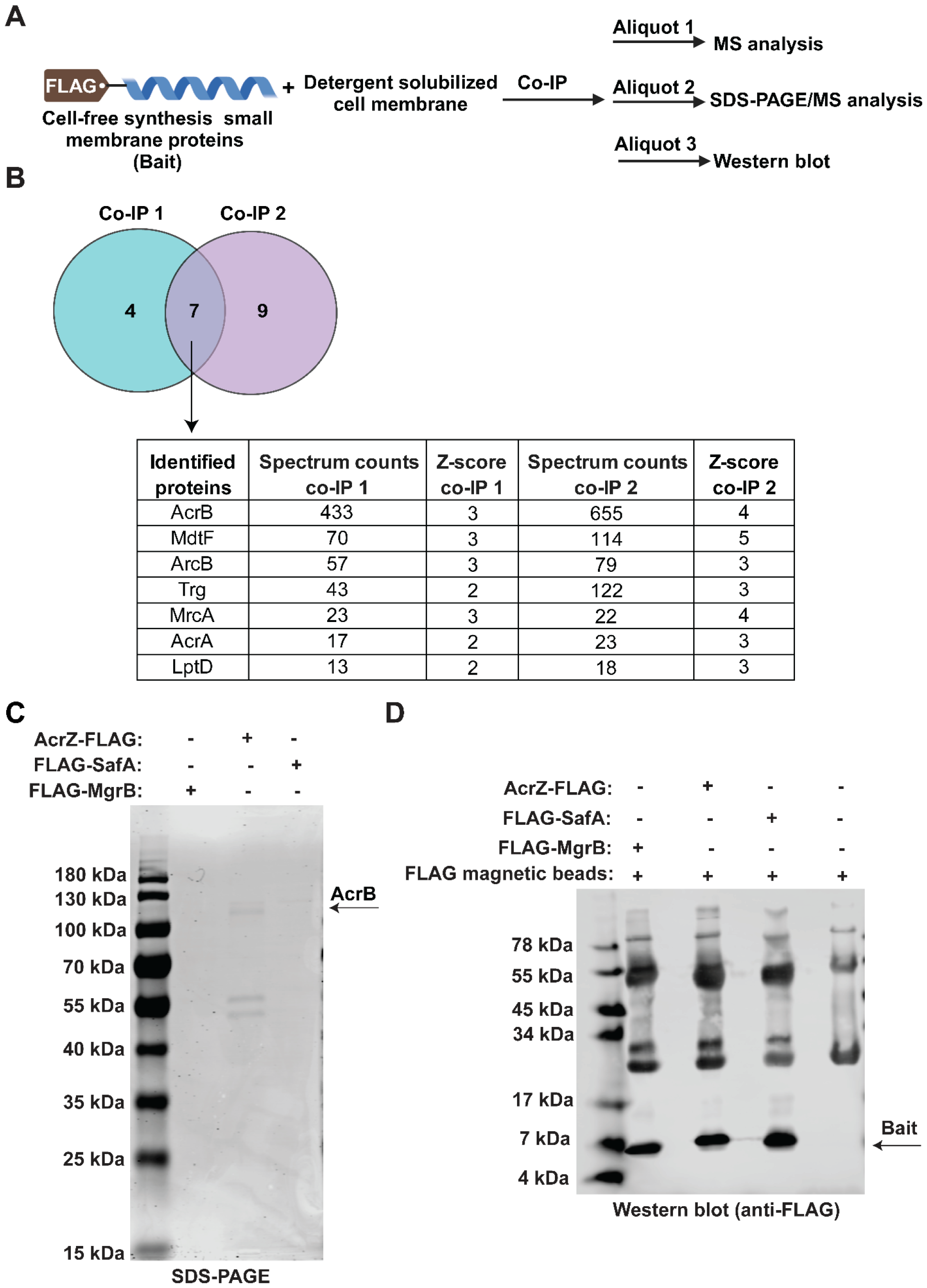
Identification of interacting targets of AcrZ. **(A)** Scheme of the *in vitro* Co-IP assay. **(B)** Using synthesized AcrZ-Flag as bait, co-immunoprecipitated proteins were identified via MS. Seven proteins with a minimum Z-score of 2 in two independent experiments are shown and considered enriched (see Materials and methods for details of Z-score calculation). The spectrum counts for each enriched protein are listed. **(C)** SDS-PAGE analysis of eluates from co-IPs using FLAG-tagged MgrB, AcrZ, and SafA as baits. **(D)** Western blot verification of synthesized FLAG-tagged MgrB, AcrZ, and SafA. Data are representative of at least two independent experiments.

Co-immunoprecipitated proteins were also eluted with 3xFLAG peptide and analyzed with SDS-PAGE (**Fig. 5C**). AcrB was identified from the top gel band with MS (**File S3**). However, PhoQ, the known interacting protein of MgrB and SafA, was not identified as one of the enriched proteins with a Z-score above 2 (**Fig. S8**), nor was it identified in the elution by SDS-PAGE (**Fig. 5C**). We did not observe any protein bands in the MgrB and SafA co-IP samples on the gel visulized with ReadyBlue® protein gel stain, even though all three bait proteins were synthesized to similar amounts in the cell-free system (**Fig. 5D**). As PhoQ activity was modulated by *in vitro* synthesized MgrB and SafA (**Fig. 3B**), we speculate that the negative co-IP results might be due to the reversible binding of the small proteins to PhoQ and the relatively low abundance of PhoQ sensor kinase in E. coli. Overall, the data indicate that our cell-free system is suitable for target discovery of small membrane proteins that form stable complexes with their targets.

## Discussion

In this study, combining lipid sponge droplets with a cell-free system, we have developed an efficient and robust approach to produce functional small membrane proteins in high yields. This allows protein function verification *in vitro* and assists function discovery of small membrane proteins by identifying their targets. The membrane-rich environment created by lipid sponge droplets provides a large membrane surface area in a three-dimensional space, supporting membrane protein synthesis and simultaneously sequestering the produced protein into the droplets. This unique feature greatly simplifies downstream experimental procedures (i.e., purification and functional assays) and reduces protein loss. Small membrane proteins sensitive to lipid composition could be synthesized in the presence of lipid sponge droplets doped with natural lipids or transferred to liposomes after their synthesis. Encouragingly, both MgrB and SafA were active without separation from lipid sponge droplets, supporting the notion that lipid sponge droplets can accommodate functional membrane proteins.

Compared to membrane protein synthesis in cell-free systems with liposomes, our system produced small membrane proteins in up to micromolar concentrations, which may be due to the unique features of lipid sponge droplets. In addition to membrane surface area, other contributing factors to *in vitro* transcription-translation may improve the protein yield of our cell-free system, including the codon usage, the ribosomal binding site (RBS) sequence, the absence of stable secondary structure of mRNA, and the optimal distance between RBS and the start codon [40]. We observed an increase in the yield of MgrB, SafA, and sarcolipin when inserting a specific DNA sequence immediately after the start codon and before the FLAG tag, which could be explained by previous reports that the sequence downstream of AUG may interact with 16S ribosomal RNA, assisting translation initiation [39]. Along the same lines, to increase the yield of small membrane proteins sensitive to N-terminal tags, one can take advantage of codon redundancy, optimizing the downstream sequence of the start codon to be more complementary to bases 1469–1483 of the 16S rRNA [39, 40]. At times, we observed minor product bands above the expected synthesized small protein in Western blots, indicating possible readthrough of the stop codon. This could likely be prevented by adding additional stop codons.

The success of using the cell-free synthesized AcrZ and identifying the known direct and indirect interactors (AcrB and AcrA) from *E. coli* total membrane highlights the system’s potential in future functional studies of small membrane proteins. Interestingly, MdtF, another multidrug efflux pump protein, was also robustly enriched via co-IP and showed good spectrum counts. MdtF is homologous to AcrB but not constitutively expressed as AcrB. The expression of MdtF was reported to be upregulated in anaerobic growth conditions in *E. coli* [41]. Its amount was expected to be less than AcrB in the total membrane isolated from *E. coli* cells grown aerobically in LB. Consistently, MS analysis of the solubilized total membrane used for the co-IP experiments indicated that the AcrB amount was about ten times higher than MdtF (data not shown). Therefore, it is likely that AcrZ interacts and possibly regulates the function of MdtF as well, though more experimental evidence is needed to support this hypothesis.

Similar to other *in vitro* systems, our cell-free approach has limitations: i) successful target discovery by co-IP appears to depend on stable protein complexes; ii) charged or polar residues within a transmembrane helix may reduce protein yield; and iii) posttranslational modifications are absent. Nevertheless, our system provides a simple protocol for small membrane protein production, functional assay, and target identification. Without cell viability requirements, it enables a high-yield production of toxic small membrane proteins or those that cause cell stress when overexpressed *in vivo*. It is versatile and supports the function of small membrane proteins. Overall, we present a robust cell-free system as an alternative approach for functional studies of small membrane proteins. With the small reaction volumes and high protein yields, our cell-free system opens up new possibilities for screens that will advance our understanding of small membrane protein biology.

## Materials and methods

### Chemical reagents and linear DNA templates

N-oleoyl β-D-galactopyranosylamine (GOA) with 99.6% purity was synthesized and analyzed by GLYCON Biochemicals. Octylphenoxypolyethoxyethanol (IGEPAL) was purchased from Sigma (I8896). The protein standard with the C-terminal FLAG tag (E-PKSH032870.10) was purchased from Biomol. Linear DNA templates were codon-optimized, produced by PCR, and purified using the GeneJET PCR purification kit (Thermo Fischer). The complete list of linear DNA templates used in this study is summarized in Table S2.

### Cell-free synthesis

The experiments were performed as described previously [31, 32]. Briefly, the PURExpress® kit (NEB) containing Solution A and Solution B was used to prepare the transcription-translation reaction. To prepare lipid sponge droplets, a lipid film was prepared by evaporating 42 µL GOA (10 mM) mixed with 100 µl chloroform in a glass vial. The lipid film was rehydrated by vortexing with 21 µL rehydration solution (14 µL PURExpress Solution A, 0.875 µL RNase Inhibitor (NEB), 17 µM IGEPAL (90mM), 3.465 µL H_2_O) to form lipid sponge droplets. To initiate protein synthesis, 3µl of the droplet solution was combined with 1.5µl PURExpress solution B and 0.5 µl DNA template solution to achieve a final concentration of 15 nM. The reactions were carried out at 37 °C at the indicated time. Lipid film and IGEPAL were omitted for cell-free synthesis without lipid sponge droplets.

### Fluorescence microscopy

A 3 µL aliquot of the cell-free reaction mix was pipetted into a lumox® dish (Sarstedt) and sealed with a cover glass. The reaction was incubated at 37 °C on stage and observed using a Nikon Eclipse Ti-E inverted fluorescence microscope with a 40× objective. The images and movies were acquired at the indicated time.

### Western blot assay

An equal volume of 2X SDS sample buffer was added to a cell-free reaction mix. The mixture was heated at 90 ⍰°C for 10 minutes and loaded onto a 10-20% SDS tris-tricine polyacrylamide gel (Thermo Fischer). Proteins were separated using electrophoresis and transferred to a polyvinylidene fluoride membrane. Anti-FLAG (Sigma) primary antibody and an IRDye 800CW-conjugated secondary antibody (LI-COR) were used to detect FLAG-tagged proteins. Protein bands were visualized using the Odyssey CLx imaging system (LI-COR) and quantified using ImageJ software.

### The overexpression and purification of PhoQ

*E.coli phoQ* with a C-terminal His_6_ tag was cloned into pET Duet-1 vector at NcoI/BamHI restriction sites and transformed into *E.coli* BL21(DE3) strain. The transformed cells were grown in the lysogeny broth (LB) medium with ampicillin (100 μg/ml) at 37 °C with vigorous shaking till OD 0.8. Protein expression was then induced with isopropyl β-D-1-thiogalactopyranoside (IPTG) at a final concentration of 0.5 mM at 16 °C overnight. Cells were harvested, resuspended in the resuspension buffer (50 mM Tris-HCl pH 8, 500 mM NaCl, 10% glycerol, and 0.1 mM PMSF), and lysed using the LM10 microfluidizer at 4 °C. After removing cell debris by centrifugation (11,000 xg) for 10 minutes, the supernatant was centrifuged again at 100,000 xg for 2 hours to pellet the membrane fraction. The pellet was then dissolved in 10 ml resuspension buffer containing 1% lauryl maltose neopentyl glycol (LMNG) (wt/vol) with gentle shaking at 4 °C for one hour. The insoluble fraction was removed by centrifugation at 21,000 xg for 45 min. The resulting supernatant was mixed with 5 ml slurry of TALON Superflow resins at 4 °C for one hour. Subsequently, the TALON Superflow resins were packed into a column and washed with 25 ml washing buffer (50 mM Tris-HCl, pH 8.0, 500 mM NaCl, 10% glycerol, 10 mM imidazole, 0.1 mM PMSF) for three times. PhoQ was eluted with 6 ml elution buffer (50 mM Tris-HCl, pH 8.0, 500 mM NaCl, 10% glycerol, 200 mM imidazole, 0.1 mM PMSF). The washing and elution buffers were not supplemented with additional LMNG due to the high concentration (1%) used in the solubilization step and the low critical micelle concentration of LMNG (0.001%). Purified PhoQ was stored in the final buffer (20 mM Tris-HCl, pH 8.0, 150 mM NaCl).

### Autophosphorylation of PhoQ in the presence of synthesized small proteins

Lipid sponge droplets containing synthesized small proteins were pelleted from a 10 µL cell-free reaction mix by centrifugation at 16,200 xg for 5 min. The supernatant was removed, followed by the addition of purified recombinant PhoQ-His_6_ in the phosphorylation buffer (50 mM Tris-HCl pH 7.5, 200 mM KCl, 0.1 mM EDTA, 10% glycerol, 0.1 mM MgSO4) at the final concentration of 14 µM, resulting in ratios of PhoQ to MgrB and SafA as 24:1 and 12:1, respectively. The autophosphorylation reaction was initiated by adding 0.1 mM ATP containing 10 μCi of [γ-^32^P]ATP and incubated at room temperature for 30 min. The reaction was stopped by adding the SDS sample buffer. The samples were then incubated at 37 °C for 2 min and subsequently loaded onto a 12% precast SDS-polyacrylamide gel (Bio-Rad). Phosphorylated PhoQ was separated from free ATPs by electrophoresis. The gel was exposed to an imaging plate overnight and analyzed with a phosphor imager (Azure Biosystems).

### In *vitro* pull-down assay

The synthesized FLAG-tagged small proteins from a 60 µL cell-free reaction mixture (approaximately 100 pmol) were used as bait for the pull-down assay. Total cell membrane was prepared from a 1L culture of *E. coli* strains (*ΔmgrB, ΔsafA, and ΔacrZ*) and solubilized in the resuspension buffer containing 1% LMNG with the final protein concentration of 8 mg/ml. The synthesized small protein was incubated with 200 µL solubilized crude membrane at 4 °C overnight with gentle rotation. After removing possible residual lipid droplets by centrifugation, anti-FLAG® M2 magnetic beads (Sigma) were added to the supernatant and incubated at 4 °C for 16 hours. The anti-FLAG® M2 magnetic beads were washed with 1 ml wash buffer (20 mM Tris-HCl, pH 8.0, 150 mM NaCl, 0.01% LMNG) for ten times and then divided into three equal aliquots: proteins retained on the beads were 1) eluted twice with 100 µl SLS at 90°C for 10min followed by MS analysis (details see below); 2) eluted with 3xFLAG peptide (Sigma) at 4 °C followed by SDS-PAGE using a precast Any kD™ polyacrylamide gel (Bio-Rad); 3) eluted with SDS sample buffer at 95 °C for 10 minutes followed by SDS-PAGE and Western blot. Proteins in 2) were visualized with Readyblue protein gel stain (Sigma). Protein bands of interest were excised from the gel and subjected to MS analysis.

### Detection of protein interactors using shotgun proteomics

The eluted proteins were reduced using 1mM TCEP at 90 °C for 10 min. Alkylation of reduced disulfide bonds was performed with 5 mM iodoacetamide at 25°C in the dark for 30 min. The SP3 approach [42] was used for further sample purification and protein digestion. Briefly, 4 µl SP3 bead slurry from the stock (20 µl beads Sera-mag beads A and B in 100 µl water) was added to the eluate. Then 500 µl acetonitrile was added, and the mixture was incubated at room temperature for 15 min. The beads were separated and washed twice with 70% ethanol, followed by washing with acetonitrile. Trypsin (1 µg) was added to the beads, and protein was digested overnight at 30°C on a shaking thermomixer. After digestion, the supernatant containing the peptides was collected. Beads were washed with water to increase peptide recovery. The total peptide pool was acidified and purified using Chromabond C18 microspin columns (Macherey-Nagel). In detail, cartridges were prepared by equilibrating with acetonitrile followed by 0.1% trifluoroacetic acid (TFA). Peptides were loaded onto equilibrated cartridges, washed with buffer containing 5% acetonitrile and 0.1% TFA and eluted with buffer containing 50% acetonitrile and 0.1% TFA. Solvent was removed, and the dried peptides were redissolved in 0.1% TFA and then analyzed using liquid-chromatography-mass spectrometry carried out on an Exploris 480 instrument connected to an Ultimate 3000 RSLC nano and a nanospray flex ion source (all Thermo Scientific).

Peptide separation was performed on a reverse phase HPLC column (75 µm x 42 cm) packed in-house with C18 resin (2.4 µm; Dr. Maisch). The following separating gradient was used: 94% solvent A (0.15% formic acid) and 6% solvent B (99.85% acetonitrile, 0.15% formic acid) to 35% solvent B over 40 minutes at a flow rate of 300 nl/min. Separated peptides were ionized at a spray voltage of 2.3 kV. The ion transfer tube temperature was set at 275 °C, and 445.12003 m/z was used as internal calibrant. The data acquisition mode was set to obtain one high-resolution MS scan at a resolution of 60,000 full width at half maximum (at m/z 200) followed by MS/MS scans of the most intense ions within 1 s (cycle 1s). To increase the efficiency of MS/MS attempts, the charged state screening modus was enabled to exclude unassigned and singly charged ions. The dynamic exclusion duration was set to 14 sec. The ion accumulation time was set to 50 ms (MS) and 50 ms at 17,500 resolution (MS/MS). The automatic gain control (AGC) was set to 3x10^6^ for MS survey scans and 2x10^5^ for MS/MS scans. The quadrupole isolation was 1.5 m/z, and the collision was induced with an HCD collision energy of 27 %.

MS raw data was then searched using Sequest HT via the Proteome Discoverer platform (Thermo Fisher Scientific) against an *E.coli* UniProt database. The search criteria were set as follows: full tryptic specificity was required (cleavage after lysine or arginine residues); three missed cleavages were allowed; carbamidomethylation (C) was set as fixed modification; oxidation (M), deamidation (N, Q) as variable modification. Mass tolerance was set to 10 ppm on precursors and 0.02 Da on fragments. The results were then imported into Scaffold (v5, Proteome Software). Within Scaffold, the data was obtained with protein FDR set to 1%, and total spectrum counts were exported for further analysis. As described before [43], we performed Z-transformation of log ratios of the spectrum counts found in AcrZ-IPs versus unrelated bait protein IP-MS experiments (MgrB-IP and SafA-IP). For calculations of spectrum log ratios, “0” was replaced with “0.5” as a background value. Proteins were considered enriched when a minimum Z-score of “2” was reached in two independently performed experiments. The Proteomics MS raw data is available via ProteomeXchange with identifier PXD051183. **Username:**, **Password:** KtWfrl3w

## Acknowledgments

We thank Roland Lill and Ulrich Mühlenhoff (Philipps University of Marburg) for support with experiments using radioactive ATPs, Jörg Kahnt (Max Planck Institute for Terrestrial Microbiology) for performing in-gel digestion and protein identification with MS, Hans-Georg Koch (University of Freiburg) for critical reading of the manuscript, members of the Yuan, the Sourjik, and the Niederholtmeyer laboratories for helpful discussions. Schematic figures were created with BioRender.com.

This work was supported by the Deutsche Forschungsgemeinschaft (DFG, German Research Foundation) priority program 2002 YU 247/3-1 and the Max Planck Society (to J.Y.). H.N. acknowledges funding by DFG grant NI 2040/1-1 and by the European Research Council (ERC starting grant SYNSEMBL, #101078028).

---

**Abstract Figure.**
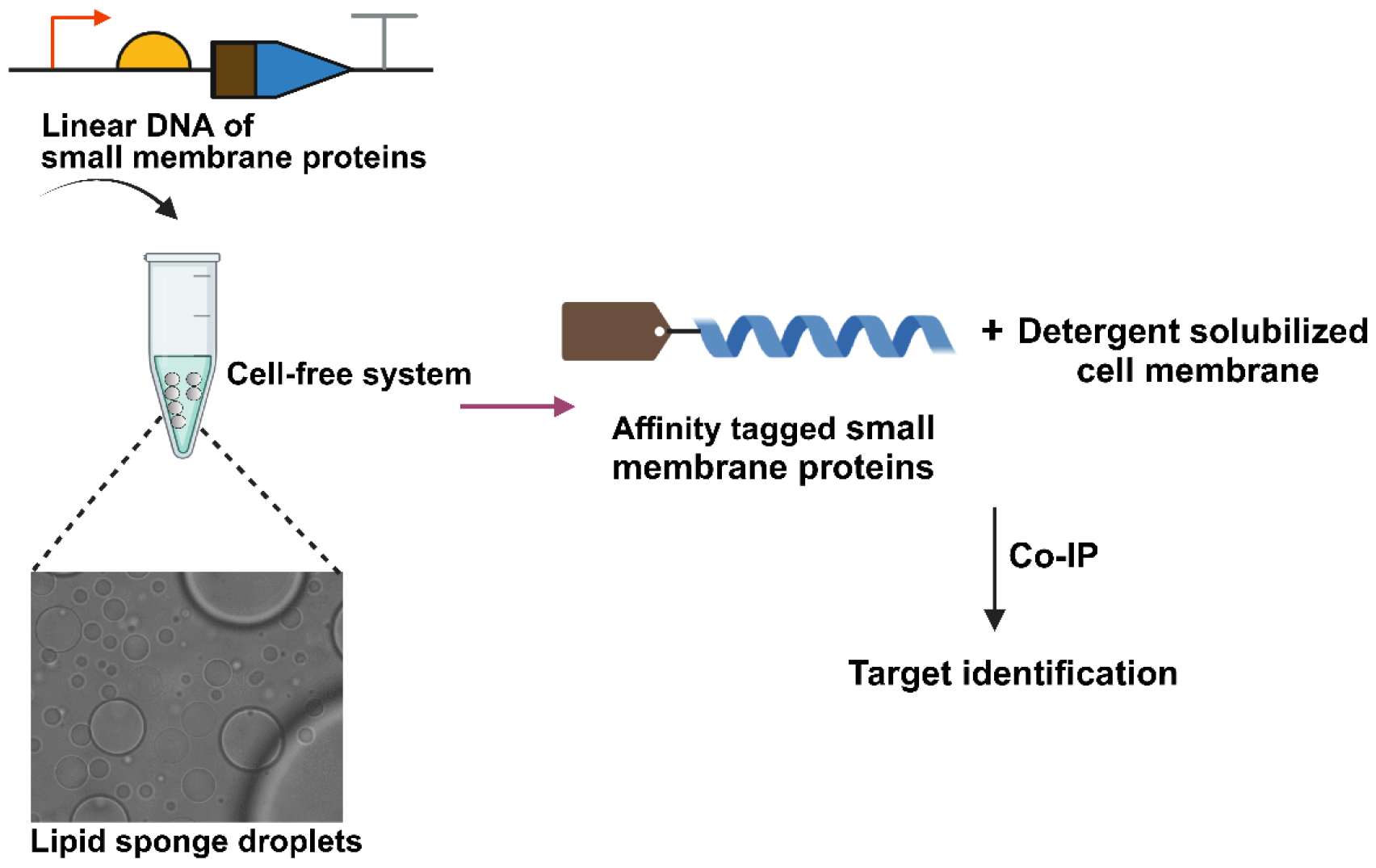
The working model involves cell-free biosynthesis of small membrane proteins in e presence of lipid sponge droplets. The figure is created with BioRender.com.

## References

1. Crook, Z.R., N.W. Nairn, and J.M. Olson, Miniproteins as a Powerful Modality in Drug Development. Trends Biochem Sci, 2020. 45(4): p. 332–346.

2. Kushwaha, A.K., et al., Plant microProteins: Small but powerful modulators of plant development. iScience, 2022. 25(11):p. 105400.

3. Feng, Y.Z., et al., Shining in the dark: the big world of small peptides in plants. Abiotech, 2023.

4. Storz, G., Y.I. Wolf, and K.S. Ramamurthi, Small proteins can no longer be ignored. Annu Rev Biochem, 2014. 83: p. 753–77.

5. Steinberg, R. and H.G. Koch, The largely unexplored biology of small proteins in pro- and eukaryotes. FEBS J, 2021. 288(24):p. 7002–7024.

6. Sberro, H., et al., Large-Scale Analyses of Human Microbiomes Reveal Thousands of Small, Novel Genes. Cell, 2019. 178(5):p. 1245–1259 e14.

7. Schlesinger, D. and S.J. Elsasser, Revisiting sORFs: overcoming challenges to identify and characterize functional microproteins. FEBS J, 2022. 289(1):p. 53–74.

8. Fesenko, I., et al., Distinct types of short open reading frames are translated in plant cells. Genome Res, 2019. 29(9):p. 1464–1477.

9. Stringer, A., et al., Identification of novel translated small ORFs in Escherichia coli using complementary ribosome profiling approaches. J Bacteriol, 2021. 204(1):p. JB0035221.

10. Duan, Y., et al., A catalogue of small proteins from the global microbiome. BioRxiv, 2023.

11. Wright, B.W., et al., The dark proteome: translation from noncanonical open reading frames. Trends Cell Biol, 2022. 32(3):p. 243–258.

12. Simoens, L., I. Fijalkowski, and P. Van Damme, Exposing the small protein load of bacterial life. FEMS Microbiol Rev, 2023. 47(6).

13. Stock, C., et al., Cryo-EM structures of KdpFABC suggest a K(+) transport mechanism via two inter-subunit half-channels. Nat Commun, 2018. 9(1):p. 4971.

14. Hobbs, E.C., et al., Conserved small protein associates with the multidrug efflux pump AcrB and differentially affects antibiotic resistance. Proc Natl Acad Sci U S A, 2012. 109(41):p. 16696–701.

15. Impens, F., et al., N-terminomics identifies Prli42 as a membrane miniprotein conserved in Firmicutes and critical for stressosome activation in Listeria monocytogenes. Nat Microbiol, 2017. 2: p. 17005.

16. Yadavalli, S.S., et al., Functional determinants of a small protein controlling a broadly conserved bacterial sensor kinase. J Bacteriol, 2020. 202(16).

17. Eguchi, Y., et al., Regulation of acid resistance by connectors of two-component signal transduction systems in Escherichia coli. J Bacteriol, 2011. 193(5):p. 1222–8.

18. Lippa, A.M. and M. Goulian, Feedback inhibition in the PhoQ/PhoP signaling system by a membrane peptide. PLoS Genet, 2009. 5(12):p. e1000788.

19. Jiang, S., et al., The inhibitory mechanism of a small protein reveals its role in antimicrobial peptide sensing. Proc Natl Acad Sci U S A, 2023. 120(41):p. e2309607120.

20. Wang, H., et al., Increasing intracellular magnesium levels with the 31-amino acid MgtS protein. Proc Natl Acad Sci U S A, 2017. 114(22):p. 5689–5694.

21. Wright, Z., et al., The small protein MntS evolved from a signal peptide and acquired a novel function regulating manganese homeostasis in Escherichia coli. Mol Microbiol, 2023.

22. Kim, E.Y., et al., Dash-and-Recruit Mechanism Drives Membrane Curvature Recognition by the Small Bacterial Protein SpoVM. Cell Syst, 2017. 5(5):p. 518–526 e3.

23. Jahn, N., S. Brantl, and H. Strahl, Against the mainstream: the membrane-associated type I toxin BsrG from Bacillus subtilis interferes with cell envelope biosynthesis without increasing membrane permeability. Mol Microbiol, 2015. 98(4):p. 651–66.

24. Savage, D.F., et al., Cell-free complements in vivo expression of the E. coli membrane proteome. Protein Sci, 2007. 16(5):p. 966–76.

25. Blanken, D., et al., Genetically controlled membrane synthesis in liposomes. Nat Commun, 2020. 11(1):p. 4317.

26. Schoborg, J.A., et al., A cell-free platform for rapid synthesis and testing of active oligosaccharyltransferases. Biotechnol Bioeng, 2018. 115(3):p. 739–750.

27. Garenne, D., et al., Cell-free gene expression. Nature Reviews Methods Primers, 2021. 1(1).

28. Jacobs, M.L., M.A. Boyd, and N.P. Kamat, Diblock copolymers enhance folding of a mechanosensitive membrane protein during cell-free expression. Proc Natl Acad Sci U S A, 2019. 116(10):p. 4031–4036.

29. Hovijitra, N.T., et al., Cell-free synthesis of functional aquaporin Z in synthetic liposomes. Biotechnol Bioeng, 2009. 104(1):p. 40–9.

30. Lyukmanova, E.N., et al., Lipid-protein nanodiscs for cell-free production of integral membrane proteins in a soluble and folded state: comparison with detergent micelles, bicelles and liposomes. Biochim Biophys Acta, 2012. 1818(3):p. 349–58.

31. Bhattacharya, A., et al., Lipid sponge droplets as programmable synthetic organelles. Proc Natl Acad Sci U S A, 2020. 117(31):p. 18206–18215.

32. Cho, C.J., et al., Functionalizing Lipid Sponge Droplets with DNA**. Chemsystemschem, 2022. 4(3).

33. Bhattacharya, A., et al., Single-Chain beta-D-Glycopyranosylamides of Unsaturated Fatty Acids: Self-Assembly Properties and Applications to Artificial Cell Development. Journal of Physical Chemistry B, 2019. 123(17):p. 3711–3720.

34. Leforestier, A., N. Lemercier, and F. Livolant, Contribution of cryoelectron microscopy of vitreous sections to the understanding of biological membrane structure. Proc Natl Acad Sci U S A, 2012. 109(23):p. 8959–64.

35. Shimizu, Y., et al., Cell-free translation reconstituted with purified components. Nature Biotechnology, 2001. 19(8):p. 751–755.

36. Kuruma, Y. and T. Ueda, The PURE system for the cell-free synthesis of membrane proteins. Nature Protocols, 2015. 10(9):p. 1328–1344.

37. Du, D., et al., Structure of the AcrAB-TolC multidrug efflux pump. Nature, 2014. 509(7501):p. 512–5.

38. Bal, N.C., et al., Sarcolipin is a newly identified regulator of muscle-based thermogenesis in mammals (vol 18, pg 1575, 2012). Nature Medicine, 2012. 18(12):p. 1857–1857.

39. Etchegaray, J.P. and M. Inouye, Translational enhancement by an element downstream of the initiation codon in Escherichia coli. J Biol Chem, 1999. 274(15):p. 10079–85.

40. Jin, X., Hong, S.H., Cell-free protein synthesis for producing ‘difficult-to-express’ proteins. Biochemical Engineering Journal, 2018. 138: p. 156–164.

41. Zhang, Y., et al., The multidrug efflux pump MdtEF protects against nitrosative damage during the anaerobic respiration in Escherichia coli. J Biol Chem, 2011. 286(30):p. 26576–84.

42. Moggridge, S., et al., Extending the Compatibility of the SP3 Paramagnetic Bead Processing Approach for Proteomics. J Proteome Res, 2018. 17(4):p. 1730–1740.

43. Ludwig, N., et al., A cell surface-exposed protein complex with an essential virulence function in Ustilago maydis. Nat Microbiol, 2021. 6(6):p. 722–730.

