## Supporting information for "A cell-free system for functional studies of small membrane proteins"

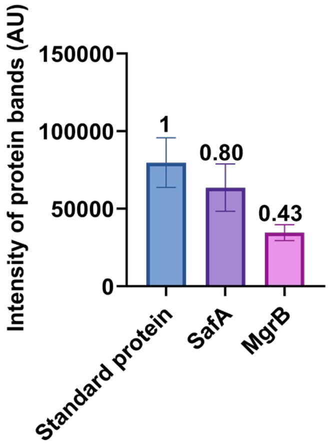


**Fig. S1. Quantification of the Western blot bands in Fig. 2B.** Compared to 0.1 µg FLAG-tagged standard protein (3.8 pmol), the amount of synthesized SafA in the presence of lipid sponge droplets in a 5µl reaction was 608 nM. The amount of synthesized MgrB reached 327 nM. The number above each bar represents the fold change of protein band intensity compared to the standard protein. ImageJ was used for quantification.

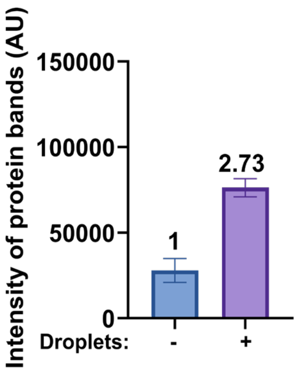


**Fig. S2. Quantification of the Western blot bands in Fig. 2C**. Compared to synthesizing SafA without droplets, the amount of synthesized SafA in the presence of lipid sponge droplets showed a 2.73-fold increase.

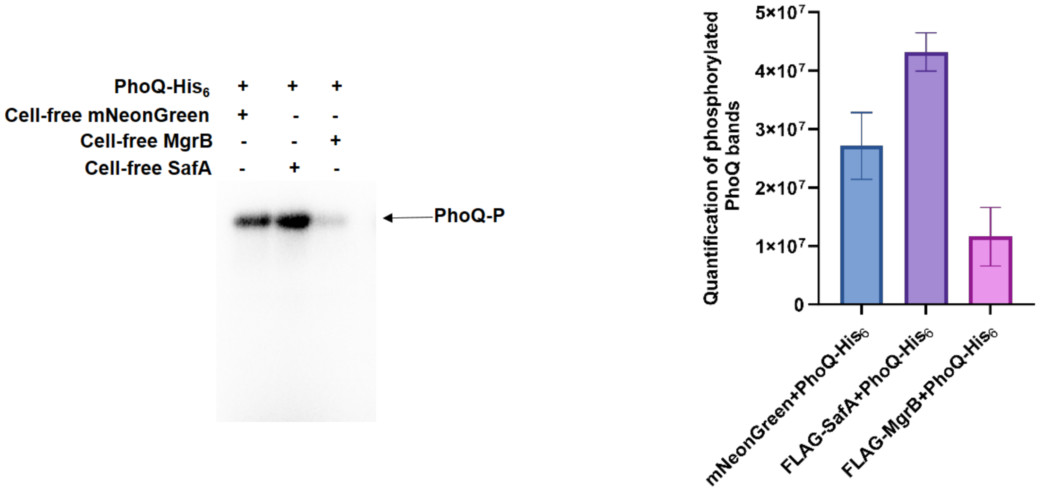


Fig. S3. **Functional assay of tag-free small membrane protein**s. The autophosphorylation reactions were performed as in Fig. 3B. Phosphorylated PhoQ in the presence of synthesized proteins were analyzed with SDS-PAGE and detected by phosphorimager (Left). Quantification of phosphorylated PhoQ bands was performed based on data from two independent experiments (Right).

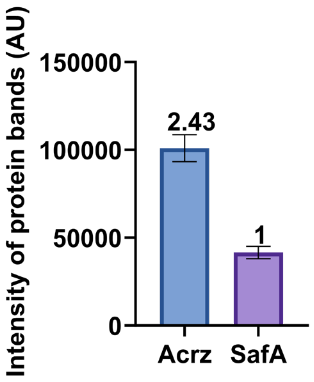


**Fig. S4. Quantification of the Western blot bands in Fig. 4A**. Compared to SafA, the amount of synthesized AcrZ showed a 2.43-fold increase (about 1.4 μM). The error bars represent standard deviations from three independent experiments.

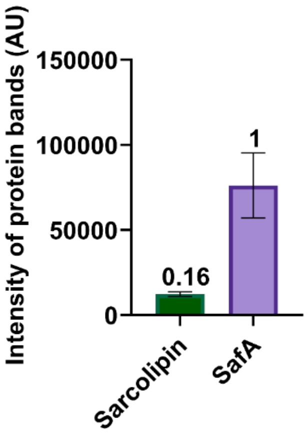


**Fig. S5. Quantification of the Western blot bands in Fig. 4B**. Compared to SafA, the amount of synthesized sarcolipin showed a 0.16-fold decrease (about 90 nM). The error bars represent standard deviations from three independent experiments.

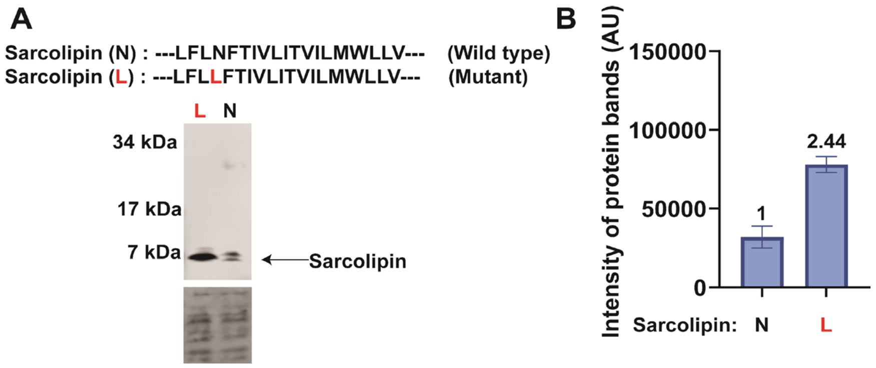


**Fig.S6. Cell-free synthesis of sarcolipin and its N to L mutant.** (A) Western blot analysis of synthesized sarcolipin and its mutant in the presence of lipid sponge droplets. The total protein stain of the PVDF membranes serves as loading control. (B) Quantification of the Western blot bands in A using ImageJ software. Compared to wild-type sarcolipin, the amount of the synthesized mutant sarcolipin showed a 2.44-fold increase. The error bars represent standard deviations from three independent experiments.

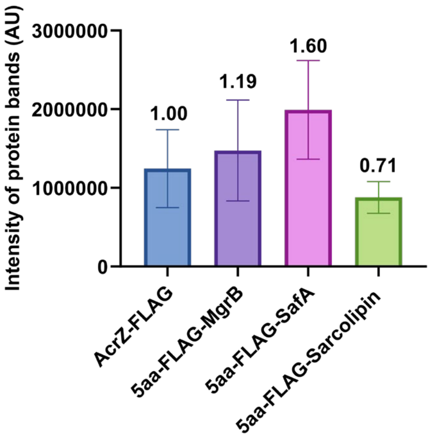


**Fig.S7. Quantification of the Western blot bands in Fig 4C.** The number above each bar represents the fold change when compared to the intensity of the AcrZ-FLAG band in the Western blot. The error bars represent standard deviations from four independent experiments.

**A**

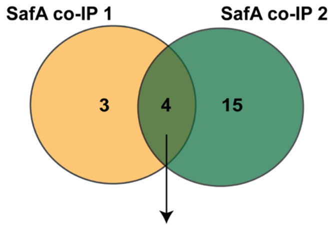


| Identified proteins | Spectrum counts-co-IP 1 | Z score-Co-IP 1 | Spectrum counts-co-IP 2 | Z score-Co-IP 2 |
|---|---|---|---|---|
| MlaF | 37 | 4 | 74 | 6 |
| MlaD | 35 | 2 | 93 | 4 |
| ZntB | 14 | 4 | 51 | 4 |
| MlaE | 3 | 2 | 27 | 5 |

**B**

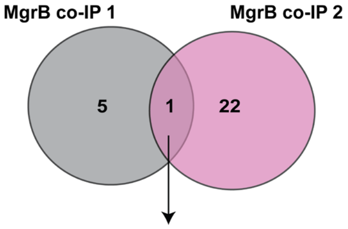


| Identified proteins | Spectrum counts-co-IP 1 | Z score-Co-IP 1 | Spectrum counts-co-IP 2 | Z score-Co-IP 2 |
|---|---|---|---|---|
| PotA | 3 | 2 | 33 | 3 |

**Fig.S8. Identification of interacting targets of MgrB and SafA.** Using synthesized FLAG-tagged SafA (A) and MgrB (B) as bait, co-immunoprecipitated proteins were identified via MS. Proteins with a minimum Z-score of 2 in two independent experiments are shown and considered enriched. The tables list enriched proteins with spectrum counts.

**TableS1**. The natural properties of small membrane proteins.

| | MgrB<br>(47 aa) | SafA<br>(65 aa) | AcrZ<br>(49 aa) | Sarcolipin<br>(31 aa) |
|---|---|---|---|---|
| **Sequences** | MKKFRWVVL<br>VVVVLACLLL<br>WAQVFNM<br>MCDQDVQFF<br>SGICAINQFIP<br>W | MHATTVKNKIT<br>QRDNYKEIMS<br>AIVVVLLLTLTLI<br>AIFSAIDQLSISE<br>MGRIARDLTHF<br>IINSLQG | MLELLKSLVFA<br>VIMVPVVMAII<br>LGLIYGLGEVFN<br>IFSGVGKKDQP<br>GQNH | MGINTRELFLN<br>FTIVLITVILMW<br>LLVRSYQY |
| **AlphaFold 2 model** | 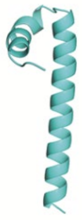 | 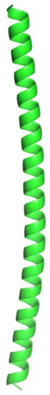 | 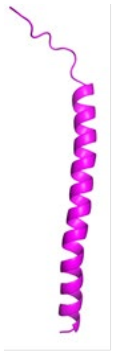 | 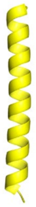 |
| **Hydrophobic ratio** | 0.57 | 0.46 | 0.51 | 0.58 |
| **Positive charge ratio** | 0.06 | 0.12 | 0.08 | 0.06 |
| **Negative charge ratio** | 0.04 | 0.08 | 0.06 | 0.03 |
| **Isoelectric point** | 7.331 | 8.137 | 6.937 | 8.347 |
| **Transmembrane helix** | WVVLVVVVL<br>ACLLLWAQV<br>F | IMSAIVVVLL<br>LTLTLIAIFSA | LVFAVIMVP<br>VVMAIILGLI<br>YG | LFLNFTIVLI<br>TVILMWLLV |

**Table S2.** The sequences of linear DNA templates were used in this study.

| **Linear DNA templates** | **Sequences**<br>**(T7 promoter, ribosomal binding site, affinity tag, ORF, T7 terminator)** |
|---|---|
| $P_{T7}$-*mNeonGreen-mgrB* | GCGAATTAATACGACTCACTATAGGCGGATAACAATTTCACACAGGAAACAG<br>ACCATGGTTTCTAAGGGTGAAGAAGACAACATGGCTTCTTTGCCAGCTACTC<br>ACGAATTGCACATCTTCGGTTCTATCAACGGTGTTGACTTCGACATGGTTGGT<br>CAAGGTACTGGTAACCCAAACGACGGTTACGAAGAATTGAACTTGAAGTCTA<br>CTAAGGGTGACTTGCAATTCTCTCCATGGATCTTGGTTCCACACATCGGTTAC<br>GGTTTCCACCAATACTTGCCATACCCAGACGGTATGTCTCCATTCCAAGCTGC<br>TATGGTTGACGGTTCTGGTTACCAAGTTCACAGAACTATGCAATTCGAAGAC<br>GGTGCTTCTTTGACTGTTAACTACAGATACACTTACGAAGGTTCTCACATCAA<br>GGGTGAAGCTCAAGTTAAGGGTACTGGTTTCCCAGCTGACGGTCCAGTTATG<br>ACTAACTCTTTGACTGCTGCTGACTGGTGTAGATCTAAGAAGACTTACCCAAA<br>CGACAAGACTATCATCTCTACTTTCAAGTGGTCTTACACTACTGGTAACGGTA<br>AGAGATACAGATCTACTGCTAGAACTACTTACACTTTCGCTAAGCCAATGGCT<br>GCTAACTACTTGAAGAACCAACCAATGTACGTTTTCAGAAAGACTGAATTGA<br>AGCACTCTAAGACTGAATTGAACTTCAAGGAATGGCAAAAGGCTTTCACTGA<br>CGTTATGGGTATGGACGAATTGTACAAGggttcttccggctcatcaggctctagtATGA<br>AAAAGTTTCGATGGGTCGTTCTGGTTGTCGTGGTGTTGGCTTGCTTGCTGCTT<br>TGGGCGCAGGTATTCAACATGATGTGCGATCAGGATGTACAATTTTTCAGCG<br>GAATTTGTGCCATTAACCAGTTTATCCCGTGGTGACTAGCATAACCCCTTGGG<br>GCCTCTAAACGGGTCTTGAGGGGTTTTTTG |
| $P_{T7}$-*mNeonGreen-safA* | GCGAATTAATACGACTCACTATAGGCGAGCTCGGTACCACAACTTAAGGAG<br>GTATTCATGGTTTCTAAGGGTGAAGAAGACAACATGGCTTCTTTGCCAGCTA<br>CTCACGAATTGCACATCTTCGGTTCTATCAACGGTGTTGACTTCGACATGGTT<br>GGTCAAGGTACTGGTAACCCAAACGACGGTTACGAAGAATTGAACTTGAAG<br>TCTACTAAGGGTGACTTGCAATTCTCTCCATGGATCTTGGTTCCACACATCGG<br>TTACGGTTTCCACCAATACTTGCCATACCCAGACGGTATGTCTCCATTCCAAG<br>CTGCTATGGTTGACGGTTCTGGTTACCAAGTTCACAGAACTATGCAATTCGA<br>AGACGGTGCTTCTTTGACTGTTAACTACAGATACACTTACGAAGGTTCTCACA<br>TCAAGGGTGAAGCTCAAGTTAAGGGTACTGGTTTCCCAGCTGACGGTCCAGT<br>TATGACTAACTCTTTGACTGCTGCTGACTGGTGTAGATCTAAGAAGACTTACC<br>CAAACGACAAGACTATCATCTCTACTTTCAAGTGGTCTTACACTACTGGTAAC<br>GGTAAGAGATACAGATCTACTGCTAGAACTACTTACACTTTCGCTAAGCCAA<br>TGGCTGCTAACTACTTGAAGAACCAACCAATGTACGTTTTCAGAAAGACTGA<br>ATTGAAGCACTCTAAGACTGAATTGAACTTCAAGGAATGGCAAAAGGCTTTC<br>ACTGACGTTATGGGTATGGACGAATTGTACAAGaagcttggctgttttggcggaggat<br>ccATGCATGCGACCACAGTGAAAAACAAAATCACGCAAAGAGACAACTATAA<br>AGAAATCATGTCTGCAATTGTGGTTGTCTTATTACTGACACTTACGTTGATAG<br>CCATTTTTTCGGCAATTGATCAGCTGAGTATTTCAGAAATGGGTCGCATTGCA<br>AGAGATCTTACACATTTCATTATCAATAGTTTGCAAGGCTGACTAGCATAACC<br>CCTTGGGGCCTCTAAACGGGTCTTGAGGGGTTTTTTG |
| $P_{T7}$-*mNeonGreen* | GTCTTCACCTCGAGGATCTTAAGGCTAGAGTAATACGACTCACTATAGGGAG<br>ATGTGGTCTAGACATTCCAGGTTAAGAAGGAGGAAAAAAAAATGGTTTCTA<br>AGGGTGAAGAAGACAACATGGCTTCTTTGCCAGCTACTCACGAATTGCACAT<br>CTTCGGTTCTATCAACGGTGTTGACTTCGACATGGTTGGTCAAGGTACTGGT<br>AACCCAAACGACGGTTACGAAGAATTGAACTTGAAGTCTACTAAGGGTGACT<br>TGCAATTCTCTCCATGGATCTTGGTTCCACACATCGGTTACGGTTTCCACCAA |

| | |
|---|---|
| | TACTTGCCATACCCAGACGGTATGTCTCCATTCCAAGCTGCTATGGTTGACGG<br>TTCTGGTTACCAAGTTCACAGAACTATGCAATTCGAAGACGGTGCTTCTTTGA<br>CTGTTAACTACAGATACACTTACGAAGGTTCTCACATCAAGGGTGAAGCTCA<br>AGTTAAGGGTACTGGTTTCCCAGCTGACGGTCCAGTTATGACTAACTCTTTGA<br>CTGCTGCTGACTGGTGTAGATCTAAGAAGACTTACCCAAACGACAAGACTAT<br>CATCTCTACTTTCAAGTGGTCTTACACTACTGGTAACGGTAAGAGATACAGAT<br>CTACTGCTAGAACTACTTACACTTTCGCTAAGCCAATGGCTGCTAACTACTTG<br>AAGAACCAACCAATGTACGTTTTCAGAAAGACTGAATTGAAGCACTCTAAGA<br>CTGAATTGAACTTCAAGGAATGGCAAAAGGCTTTCACTGACGTTATGGGTAT<br>GGACGAATTGTACAAGTAACGACTCAGGCTGCTACTCAAAACTAGCATAACC<br>CCTTGGGGCCTCTAAACGGGTCTTGAGGGGTTTTTTG |
| $P_{T7}$-*flag-mgrB* | GCGAATTAATACGACTCACTATAGGAATTGTGAGCGGATAACAATTTCACAC<br>AGGAAACAGACCATGGATTATAAAGATGATGATGATAAAggtggcggaggatcc<br>ATGAAAAAGTTTCGATGGGTCGTTCTGGTTGTCGTGGTGTTGGCTTGCTTGC<br>TGCTTTGGGCGCAGGTATTCAACATGATGTGCGATCAGGATGTACAATTTTT<br>CAGCGGAATTTGTGCCATTAACCAGTTTATCCCGTGGTGACTAGCATAACCCC<br>TTGGGGCCTCTAAACGGGTCTTGAGGGGTTTTTTG |
| $P_{T7}$- *flag-safA* | GCGAATTAATACGACTCACTATAGGTCTCCATACCCGTTTTTTTGGGCTAGCG<br>AGGAGTTCGAGCTCATGGATTATAAAGATGATGATGATAAAggtggcggtggca<br>gcATGCATGCGACCACAGTGAAAAACAAAATCACGCAAAGAGACAACTATAA<br>AGAAATCATGTCTGCAATTGTGGTTGTCTTATTACTGACACTTACGTTGATAG<br>CCATTTTTTCGGCAATTGATCAGCTGAGTATTTCAGAAATGGGTCGCATTGCA<br>AGAGATCTTACACATTTCATTATCAATAGTTTGCAAGGCTGACTAGCATAACC<br>CCTTGGGGCCTCTAAACGGGTCTTGAGGGGTTTTTTG |
| $P_{T7}$-*mgrB* | GCGAATTAATACGACTCACTATAGGTTTTTTTGGGCTAGCGAGGAGTTCGAG<br>CTCATGAAAAAGTTTCGATGGGTCGTTCTGGTTGTCGTGGTGTTGGCTTGCTT<br>GCTGCTTTGGGCGCAGGTATTCAACATGATGTGCGATCAGGATGTACAATTT<br>TTCAGCGGAATTTGTGCCATTAACCAGTTTATCCCGTGGTGACTAGCATAACC<br>CCTTGGGGCCTCTAAACGGGTCTTGAGGGGTTTTTTG |
| $P_{T7}$- *safA* | GCGAATTAATACGACTCACTATAGGCGGATAACAATTTCACACAGGAAACAG<br>ACCATGCATGCGACCACAGTGAAAAACAAAATCACGCAAAGAGACAACTAT<br>AAAGAAATCATGTCTGCAATTGTGGTTGTCTTATTACTGACACTTACGTTGAT<br>AGCCATTTTTTCGGCAATTGATCAGCTGAGTATTTCAGAAATGGGTCGCATTG<br>CAAGAGATCTTACACATTTCATTATCAATAGTTTGCAAGGCTGACTAGCATAA<br>CCCCTTGGGGCCTCTAAACGGGTCTTGAGGGGTTTTTTG |
| $P_{T7}$-*acrZ-flag* | GAAATTAATACGACTCACTATAGGGGAATTGTGAGCGGATAACAATTCCCCT<br>GTAGAAATAATTTTGTTTAACTTTAATAAGGAGATATACCATGGGCTTAGAG<br>TTATTAAAAAGTCTGGTATTCGCCGTAATCATGGTACCTGTCGTGATGGCCAT<br>CATCCTGGGTCTGATTTACGGTCTTGGTGAAGTATTCAACATCTTTTCTGGTG<br>TTGGTAAAAAAGACCAGCCCGGACAAAATCATggcggtggcggtagcGATTATAA<br>AGATGATGATGATAAATGACTAGCATAACCCCTTGGGGCCTCTAAACGGGTC<br>TTGAGGGGTTTTTTG |
| $P_{T7}$- *flag-sacrcolipin* | GCGAATTAATACGACTCACTATAGGAATTGTGAGCGGATAACAATTTCACAC<br>AGGAAACAGACCATGGATTATAAAGATGATGATGATAAAggtggcggaggatcc<br>ATGGGCATTAACACCCGTGAGCTGTTTCTGAACTTCACTATTGTCTTGATTAC |

| | |
|---|---|
| | GGTTATTCTTATGTGGCTCCTTGTGCGTTCCTATCAGTACTGACTAGCATAAC<br>CCCTTGGGGCCTCTAAACGGGTCTTGAGGGGTTTTTTG |
| $P_{T7}$- *flag-sacrcolipin*(N11L) | GCGAATTAATACGACTCACTATAGGAATTGTGAGCGGATAACAATTTCACAC<br>AGGAAACAGACCATGGATTATAAAGATGATGATGATAAAggtggcggaggatcc<br>ATGGGCATTAACACCCGTGAGCTGTTTCTGCTGTTCACTATTGTCTTGATTAC<br>GGTTATTCTTATGTGGCTCCTTGTGCGTTCCTATCAGTACTGACTAGCATAAC<br>CCCTTGGGGCCTCTAAACGGGTCTTGAGGGGTTTTTTG |
| $P_{T7}$-5aa -*flag-safA* | GAAATTAATACGACTCACTATAGGGGAATTGTGAGCGGATAACAATTCCCCT<br>GTAGAAATAATTTTGTTTAACTTTAATAAGGAGATATACCATGGGCTTAGAG<br>TTAGATTATAAAGATGATGATGATAAAggtggcggtggcagcATGCATGCGACCA<br>CAGTGAAAAACAAAATCACGCAAAGAGACAACTATAAAGAAATCATGTCTG<br>CAATTGTGGTTGTCTTATTACTGACACTTACGTTGATAGCCATTTTTTCGGCA<br>ATTGATCAGCTGAGTATTTCAGAAATGGGTCGCATTGCAAGAGATCTTACAC<br>ATTTCATTATCAATAGTTTGCAAGGCTGACTAGCATAACCCCTTGGGGCCTCT<br>AAACGGGTCTTGAGGGGTTTTTTG |
| $P_{T7}$-5aa-*flag-sarcolipin* | GAAATTAATACGACTCACTATAGGGGAATTGTGAGCGGATAACAATTCCCCT<br>GTAGAAATAATTTTGTTTAACTTTAATAAGGAGATATACCATGGGCTTAGAG<br>TTAGATTATAAAGATGATGATGATAAAggtggcggaggatccATGGGCATTAACA<br>CCCGTGAGCTGTTTCTGAACTTCACTATTGTCTTGATTACGGTTATTCTTATGT<br>GGCTCCTTGTGCGTTCCTATCAGTACTGACTAGCATAACCCCTTGGGGCCTCT<br>AAACGGGTCTTGAGGGGTTTTTTG |
| $P_{T7}$-5aa-*flag-mgrB* | GAAATTAATACGACTCACTATAGGGGAATTGTGAGCGGATAACAATTCCCCT<br>GTAGAAATAATTTTGTTTAACTTTAATAAGGAGATATACCATGGGCTTAGAG<br>TTAGATTATAAAGATGATGATGATAAAggtggcggaggatccATGAAAAAGTTTC<br>GATGGGTCGTTCTGGTTGTCGTGGTGTTGGCTTGCTTGCTGCTTTGGGCGCA<br>GGTATTCAACATGATGTGCGATCAGGATGTACAATTTTTCAGCGGAATTTGT<br>GCCATTAACCAGTTTATCCCGTGGTGACTAGCATAACCCCTTGGGGCCTCTAA<br>ACGGGTCTTGAGGGGTTTTTTG |

**Movie S1.** Cell-free synthesis of mNeonGreen-MgrB in the presence of lipid sponge droplets shown as a time-lapse movie of merged transmitted light and GFP fluorescence images. The movie is associated with the images in Figure 1A.
